# Activation of the endoplasmic reticulum unfolded protein response reverses an inflammation-like response to cytoplasmic DNA

**DOI:** 10.1101/440883

**Authors:** Ashley B. Williams, Felix Heider, Jan-Erik Messling, Wilhelm Bloch, Björn Schumacher

## Abstract

Innate immune responses protect organisms against various insults, but may lead to tissue damage when aberrantly activated. In higher organisms, cytoplasmic DNA can trigger inflammation that can lead to tissue degeneration. Simpler *in vivo* models could shed new mechanistic light on how inflammatory responses to cytoplasmic DNA lead to pathology. Here we show that in DNase II-deficient *Caenorhabditis elegans*, persistent foreign cytoplasmic DNA leads to systemic tissue degeneration, and we identify impaired protein homeostasis as an inflammatory pathomechanism. This pathological outcome can be alleviated by improving protein homeostasis, either via ectopic induction of the endoplasmic reticulum unfolded protein response (UPR^ER^) or by treatment with *N*-acetylglucosamine. Our results establish *C. elegans* as an ancestral metazoan model for studying outcomes of inflammation-like conditions caused by persistent cytoplasmic DNA and provide insight into potential therapies for conditions involving chronic inflammation.

## INTRODUCTION

The processes of sensing and responding to foreign DNA are important in a variety of contexts for various organisms, from bacteria to humans. In the perhaps the simplest example, bacteria possess at least two mechanisms for responding to foreign DNA: restriction-modification systems (Vasu and Nagaraja, 2013), which protect the bacteria from foreign DNA (especially during bacteriophage infection), and the CRISPR-Cas9 system, which functions as a rudimentary adaptive immune response to foreign DNA (Ishino et al., 2018). Much more complex systems have evolved in higher eukaryotes (Dhanwani et al., 2018; Gallucci and Maffei, 2017; Gasser et al., 2017). In mice and humans, a complex array of detection systems has been identified that are deployed to sense and respond to inappropriately-localized DNA. Well-characterized DNA sensing pathways include Toll-like receptor-mediated sensing in endosomes (via TLR9), and direct sensing in the cytoplasm by cGAS-STING-IRF3 pathway and the AIM2 inflammasome pathway. While these systems serve important functions in protecting the host organism against bacteria and viruses, failures in various cellular processes that lead to chronic immune signaling, for instance to host-derived DNA, can effectively turn these normally protective pathways against the host (Dhanwani et al., 2018).

TLR9 sensing of self-DNA can lead to detrimental effects due to improper activation of inflammatory pathways, which under certain circumstances can become chronic. For example, in mice, the injection of fetal DNA, which is detected by TLR9 due to its hypomethylation, can lead to complications and fetal loss via an inappropriate inflammatory response to the DNA (Scharfe-Nugent et al., 2012). When self-derived DNA is not degraded due to defects in DNase II-dependent degradation, several inflammation-associated phenotypes have been described. For example, DNase II-deficient mice show a polyarthritis-like disease that affects the joints due to the improper localization of host DNA in phagocytic cells, where it can then be detected by AIM2 to stimulate an autoinflammatory condition that subsequently leads to joint degeneration (Jakobs et al., 2015). A similar polyarthritis-like condition has been demonstrated in DNase-II deficient mice due to a failure in the degradation of self-chromosomal DNA during erythropoiesis, which leads to chronic TNF-alpha production and joint degeneration (Kawane et al., 2006). Similar outcomes have been reported in mice after injection of mitochondrial or bacterial DNA directly into the joints (Collins et al., 2004). During autophagy of damaged mitochondria, DNase II is responsible for degrading the residual mitochondrial DNA (Kawane et al., 2006). Because of the similarities between mitochondrial and bacteria DNA, especially the lack of methylation, mitochondrial DNA can be recognized by host DNA sensing pathways, in particular TLR9. In DNase II-deficient mice, such a lack of DNA degradation during autophagy of mitochondria in cardiomyocytes leads to a cell-autonomous inflammatory response that can ultimately lead to heart failure (Oka et al., 2012).

The common feature of these processes is a profound loss of tissue integrity and functionality that are generally ascribed to induction of a chronic innate immune response. What is still lacking is a detailed understanding the sub-cellular and molecular transactions that lead to these phenotypic outcomes. In an attempt to explore these unresolved questions in a simpler metazoan model, we took advantage of advances in handling and manipulation methods in *Caenorhabditis elegans* that allowed us to directly inject purified DNA into the cytoplasm of the intestinal cells of this one-millimeter-long animal. Using this approach, we could study the resulting effects on tissue integrity. To expand our analysis of the effects of foreign DNA on tissues in a situation more likely to be encountered under environmental conditions, we developed a method to deliver foreign DNA into the cytoplasm of the worm’s intestinal cells using infection with a pathogenic bacterial strain of human origin.

No DNA-sensing pathways have been discovered in *C. elegans;* however, they do possess a rudimentary, transcription-based innate immune response (Ermolaeva and Schumacher, 2014). Several innate immunity-related regulators have been identified—the best characterized factors of FSHR-1(Powell et al., 2009) and PMK-1(Kim et al., 2002), the latter of which is the nematode homolog of the human p38 MAP kinase. The primary outcome of the activation of these pathways is a robust transcriptional induction of many diverse genes (Shivers et al., 2008), most of whose functions have yet to be characterized. While it is clear that these pathways confer resistance to a broad range of pathogens, the mechanisms remain incompletely understood. What is becoming clear is that, like in the case of the transcriptional responses to induction of the human innate immune response, the activation of these pathways comes with a trade-off between the beneficial effects in imparting pathogen resistance and organismal stress due to the burden of the massive induction in gene expression (Cheesman et al., 2016; Head et al., 2017).

Through our analysis, we demonstrate that *C. elegans* responds to foreign DNA, and that the outcome of persistent, foreign, cytoplasmic DNA is a systemic decline in tissue integrity and functionality. We characterized the subcellular and molecular mechanism of this phenotype, and we describe here novel therapeutic interventions that can alleviate these detrimental effects. Our results may have implications in understanding and treating similar pathological outcomes in humans.

## RESULTS

### *C. elegans* is sensitive to persistent cytoplasmic DNA

To develop *C. elegans* as a model to study organismal effects of foreign DNA, we devised a method for the direct injection of foreign DNA into the cytoplasm of intestinal cells. These cells for chosen for two reasons: (1) from a practical standpoint, the cells are large and facilitate reproducible injection; (2) the intestine seems to be a center of immune function in the worm (Ermolaeva and Schumacher, 2014; Kim and Ewbank, 2018; Pukkila-Worley and Ausubel, 2012; Yuen and Ausubel, 2018) and, therefore, its constituent cells are good candidates for responding to foreign DNA. The intestine consists of a ring of four cells, followed by 8 rings formed by two cells each, which combined for a tube structure with a central lumen. To maximize reproducibility, we first injected purified *E. coli* genomic DNA into one cell of the second intestinal ring using an Axio Observer A1 (Zeiss) and FemtoJet (Eppendorf) equipped with Femtotip II needles (Figure 1A) (for detailed methods, see *Materials and Methods*). To monitor the location and success of the injection, the fluorescent compound rhodamine-isothiocyanate dextran (10,000 MW) was included in the injection mixture, as it is commonly used for this purpose since it is generally inert in cells and has limited ability to diffuse (Schmued et al., 1990).

**Figure 1.**
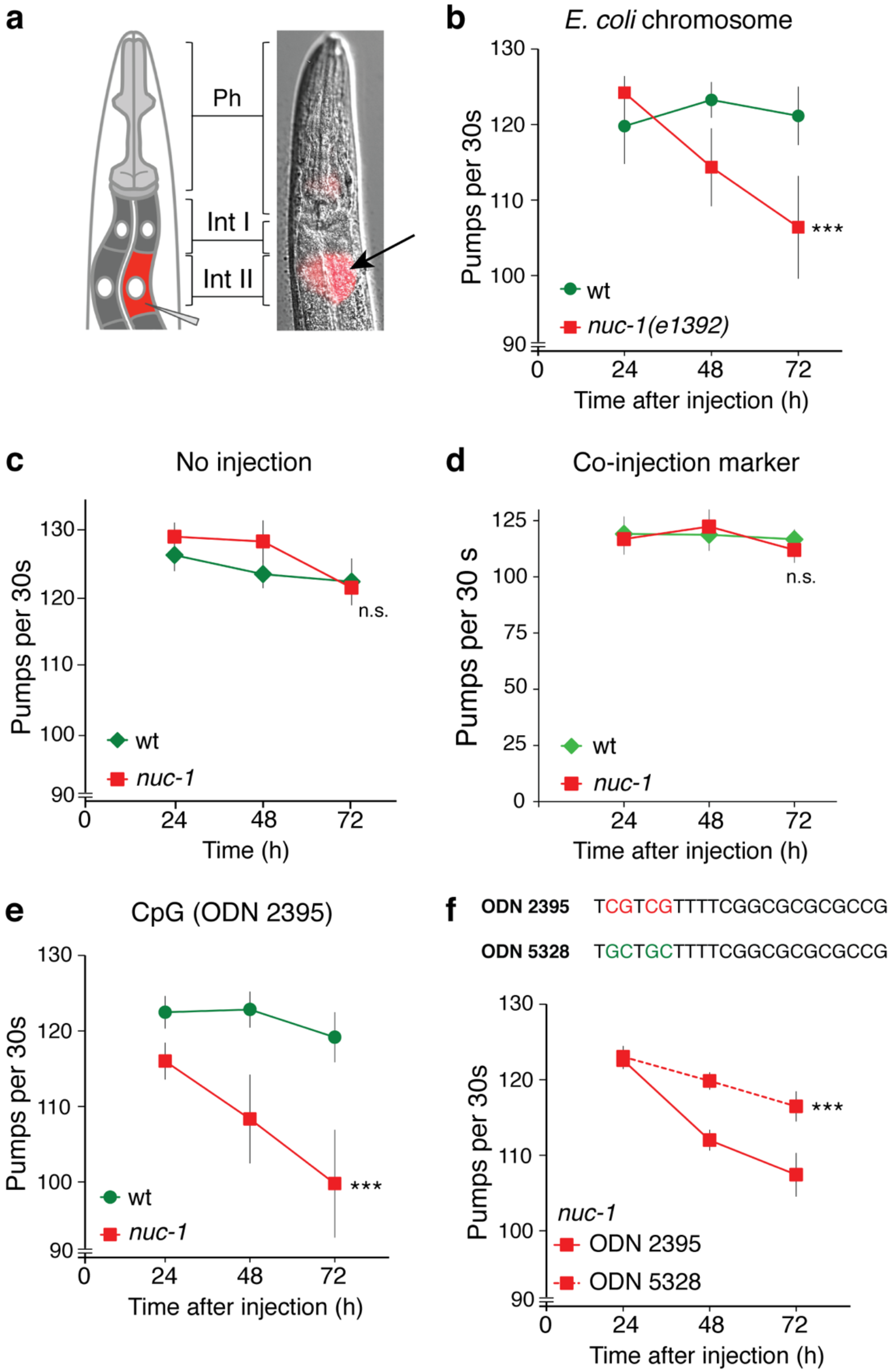
Injection of DNA into intestinal cells causes declines in tissue functionality in *nuc-1* worms. **a.** A cartoon showing the approach to intestinal injection. Ph, pharynx; Int I, Int II, intestinal rings I and II; arrow indicates the approximate location of the injection. **b.** Plot of the pharyngeal pumping rates of wt and *nuc-1* worms injected with purified ECO85 genomic DNA. **c.** Plot of the pharyngeal pumping rates of uninjected wt and *nuc-1* worms over the first 72 hours after reaching the young adult stage (24 h after L4). **d.** Plot of the pharyngeal pumping rates of wt and *nuc-1* worms injected the fluorescent co-injection marker in the same buffer (rhodamine-dextran) without DNA. **c.** Plot of the pharyngeal pumping rates of uninjected wt and *nuc-1* worms over the first 72 hours after reaching the young adult stage (24 h after L4). **d.** Plot of the pharyngeal pumping rates of the same strains injected with synthetic CpG-containing oligoribodeoxynucleotide (ODN 2395). **f.** (**top**) Comparison of the sequences of ODN 2395 and the control, ODN 5328. Red bases show the CpG sequences in ODN 2395 and blue bases highlight the same positions in the control ODN 5328. (**bottom**) Plot of the pharyngeal pumping rates of *nuc-1* worms injected with ODN 2395 or ODN 5328. Each plot shows the mean ± S.D at each time point. The statistical significance of the differences in the trends between the control and experimental groups was assessed using two-way ANOVA (n.s. P > 0.05, ** 0.01 > P > 0.001, *** P < 0.001).

To monitor consequences of DNA injections on the overall health of the animals we measured pharyngeal pumping, which is an established and highly sensitive method for monitoring even small changes in tissue functionally during aging (Bolanowski et al., 1981; Chow et al., 2006; Huang et al., 2004), following induction of DNA damage (Wilson et al., 2017), and during infection (O’Quinn et al., 2001; Tan et al., 1999). DNase II-defective worms have been previously shown to induce a mild immune response, even in the absence of a pathogen (Yu et al., 2015); therefore, we suspected that a DNase II-mutant strain (in this case carrying the *nuc-1(e1392)* allele(Wu et al., 2000)) may be a useful tool for our experiments. In wildtype worms, we did not observe any change in the pumping rate following injection of *E. coli* genomic DNA; however, identical injection into DNase II-deficient worms, in which the DNA should persist, led to a statistically significant decline in the pumping rate over the course of 72 hours post-injection (Figure 1B). Importantly, such a difference in pumping rates was not observed in either un-injected animals or animals injected with the co-injection marker alone (Figure 1C and 1D).

Injection of whole *E. coli* genomic DNA proved technically challenging, as its large molecular weight tended to clog the microinjection needles; therefore, we switched to low molecular weight CpG-rich synthetic oligonucleotides (ODNs). As shown in Figure 1E, injection of a CpG-containing ODN (ODN 2395) led to a similar pumping rate decline. In contrast to the results obtained with *E. coli* genomic DNA, a slight reduction of the pumping rate of wildtype worms resulted after ODN injection. We attribute this difference to increased resistance of the ODNs to DNase digestion due to phosphorothioate linkages in the ODN backbone.

A common feature shared between the *E. coli* genomic DNA and the synthetic ODNs is the presence of CpG sites. In higher organisms, CpG-containing DNA introduced by bacterial pathogens can be sensed by the toll-like receptor (TLR) 9 to induce an immune response. We supposed that a similar process might occur in *C. elegans;* to test this hypothesis, we compared the effects of a CpG-containing ODN (as in Figure 1E) to non-CpG control ODNs (ODN 5328) (which are non-antagonistic for mammalian TLR9) (Figure 1F, top). As shown in Figure 1F (bottom), injection of the control ODN caused a milder decrease in pumping rate, suggesting that worms have a more robust response to CpG-containing DNA. This finding makes an important connection between the responses of worms and higher organisms to foreign DNA.

### Cytoplasmic DNA accumulates in intestinal cells during infection with uropathogenic *E. coli* (UPEC) and results in a systemic health decline

A limitation of our injection experiments was that we could introduce only small amounts of foreign DNA into the intestinal cells, likely preventing the detection of more robust effects on the animals that could result from higher loads of cytoplasmic DNA; therefore, a means of introducing larger amounts of DNA could be useful. Some pathogenic *E. coli*, including some UPEC strains, can invade host cells (Lewis et al., 2016); therefore, we predicted that infection of the intestinal lumen of a DNase II-defective mutant worm strain with UPEC might provide such a possibility. We obtained a clinical UPEC isolate (designated ECO85) and infected worms via feeding. To determine if bacterial DNA accumulated in the cytoplasmic of UPEC-infected worms, we stained dissected intestinal cells with 4′,6-diamidino-2-phenylindole (DAPI) to examine the DNA content of the cells. Confocal imaging (Figure 2A) of stained animals revealed that, in addition to the bacterial DNA in the intestinal lumen of both wildtype and DNase II-defective worms (outlined with yellow dashed lines), DAPI-stained bodies similar in size to the luminal DNA bacterial DNA bodies accumulated preferentially in the cytoplasm of the DNase II-defective worms (examples indicated by white arrow heads) (Figures 2A-C). The observation that both wildtype and DNase II-deficient worms contain such bodies suggests that the bacterial DNA is introduced into the cells of both worm strains, but that it persists more robustly in the DNase II-defective strain. Notably, similar DAPI-stained bodies were not observed in DNase II-deficient animals grown on the non-pathogenic *E. coli* strain OP50 (Supplemental Figure 1). Interesting, we were not able to identify any other bacterial components in the cytoplasm of the intestinal cells using anti-*E. coli* antibodies or transmission electron microscopy (TEM), arguing against direct cellular invasion; however, it is now clear that bacterial DNA can enter cells via outer membrane vesicles (OMVs) and that is persists (Bitto et al., 2017). Therefore, we propose that the bacterial DNA may enter the cells as cargo in OMVs, although this question remains to be fully addressed. We were next curious if the persistent foreign DNA introduced via UPEC infection may lead to preferential declines in tissue integrity and functionality in the DNase II-deficient worms compared to wildtype worms. To address this question, we looked at several indicators of health in the worms: (1) pharyngeal pumping rate; (2) intestinal tissue integrity; (3) muscle integrity; and (4) reproductive function.

**Figure 2.**
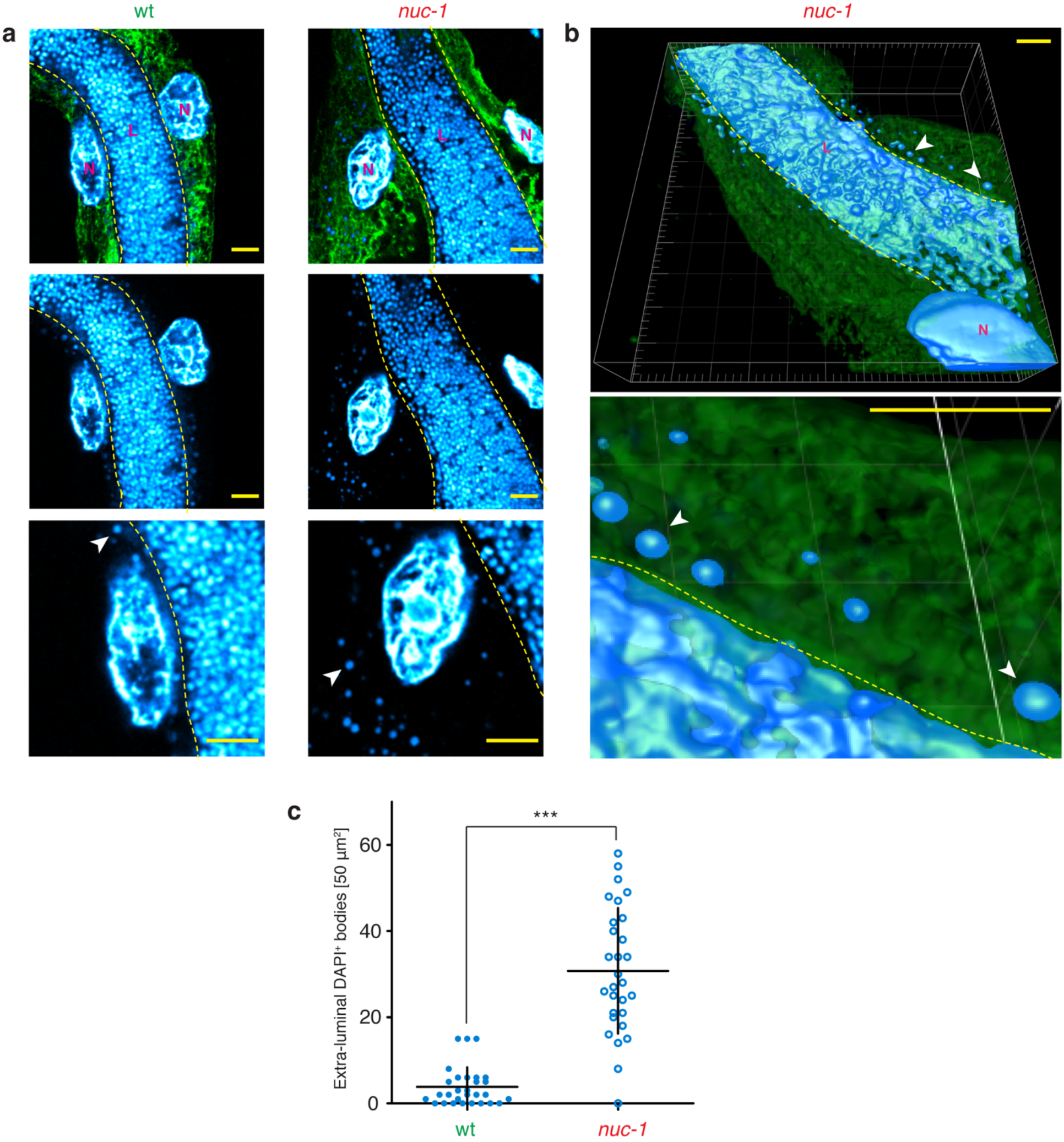
Cytoplasmic DNA preferentially accumulates in the cytoplasm of intestinal cells in UPEC-infection *nuc-1* worms. a. *top panels:* single confocal Z-planes from ECO85-infected wt and *nuc-1* worms showing two intestinal cells. ***middle panels:*** only the DAPI (DNA) signal is shown. ***bottom panels:*** enlarged images of the DAPI channel of the same images. Green indicates α-tubulin and cyan indicates DAPI; N, nucleus; L, intestinal lumen; yellow line marks the margin between the lumen and intestinal cells; arrows point to representative DAPI-staining inclusions. For all images, the scale bars represent 5 µm. **b.** Three-dimension reconstructions from confocal images showing the positions of the DAPI-staining bodies within the cytoplasm of the intestine. **c.** Quantification of the number of DAPI-staining inclusions in ECO85-infected wt and *nuc-1* worms. All samples are shown with the means ± S.D. Statistical significance was assessed using a unpaired, two-tailed Student’s *t* test (*** P < 0.001).

The pharyngeal pumping rate was measured (as above) during the first five days after UPEC infection. As shown in Figure 3A, UPEC infection led to a more rapid decline in the pumping rate in wildtype and DNase II-deficient worms compared to animals reared on *E. coli* OP50 (the normal food source for *C. elegans*); however, this effect was much more severe in the DNase II-deficient animals. We confirmed this effect with two alleles of *nuc-1: e1392* and *n887.* The variation between the phenotypes following injection and infection could be due to differences in the amount of cytoplasmic DNA delivered into the intestinal cells in each technique, as injected worms – which receive much less DNA – showed a smaller pumping rate decline (Figure 1).

**Figure 3.**
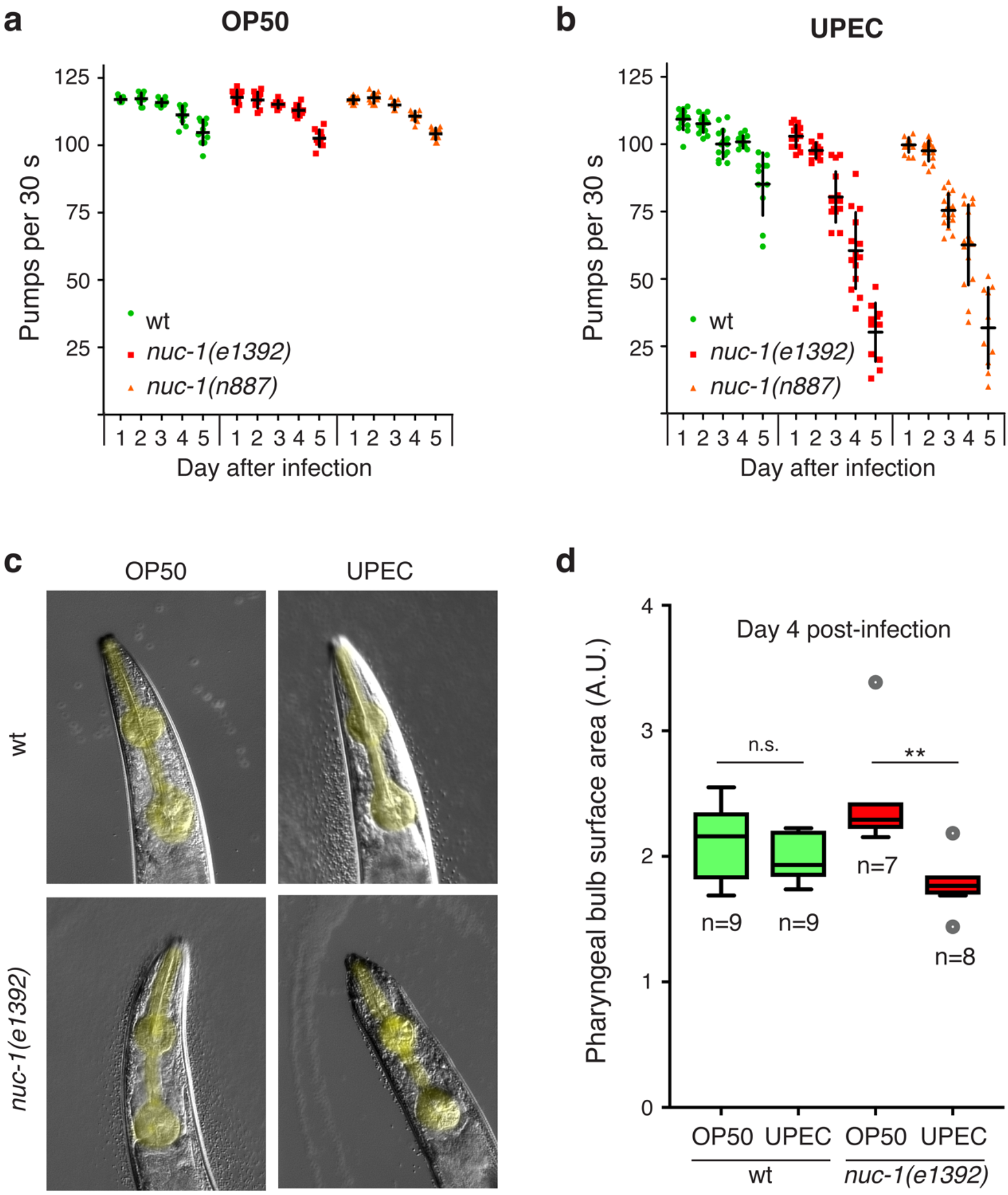
UPEC infection induces a functional decline in the pharynx. Pharyngeal pumping rates of OP50-mock infected (**a**) and UPEC-infected (**b**) wt and *nuc-1* worms. Measurements for each individual worm are shown with the mean ± S.D. **c.** Representative differential interference contrast (DIC) images of pharynxes from OP50-mock infected and UPEC-infected wt and *nuc-1* worms on day 4 (corresponding to day 4 in a and b). The pharynxes are highlighted in yellow. **c.** Quantification of the combined surface areas of the pharyngeal metacorpus and posterior bulb in OP50-mock infected and UPEC-infected wt and *nuc-1* worms on day 4. Tukey plots are used to show the interquartile range and the median (see the *Experimental Procedures* and Supplemental Figure 1). Statistical significance was assessed using unpaired, two-tailed Student’s *t* tests; ** 0.01 > P > 0.001, n.s. indicates no significant difference.

Changes in pharyngeal pumping in *C. elegans* can be attributed either to behavioral changes (reviewed in (Luedtke et al., 2010)) or to loss of tissue functionality. The validity of our pumping assay to address our experimental questions depends on the latter possibility; therefore, we wanted to confirm that the integrity of the pharynx was preferentially compromised in the UPEC-infected DNase II-defective worms. We imaged worms using differential interference contrast (DIC) microscopy four days after infection (example images are provided in Figure 3c) and then measured the surface areas of the pharyngeal metacorpus and posterior bulb in each worm (explained in more detail in Supplemental Figure 1). These values were added and plotted in arbitrary units as shown in Figure 3d. Indeed, we observed a pronounced and statistically significant decrease in surface area in the *nuc-1* mutant animals fed UPEC compared to the *nuc-1* animals fed OP50. No significant difference was observed under the same conditions in the wild type worms. These results demonstrate that the pumping rate declines in our experimental system are the result of pharyngeal atrophy and not behavioral changes; therefore, this assay provides a reliable and quantitative metric for tissue functionality.

Figure 4 shows images of the anterior sections of intestines (adjacent to the pharynx) in animals stained with the dyes acridine orange (AO) or Nile red (NR), both of which accumulate in punctate intracellular acidic compartments (Clokey and Jacobson, 1986; O’Rourke et al., 2009). Dissolution of the punctate bodies is a hallmark of necrotic cell death and can be used as a marker for tissue integrity during pathogen infection (Zou et al., 2014). The intestines of wildtype and DNase II-deficient worms grown on *E. coli* OP50 show accumulations of AO and NR in highly resolved punctate bodies, as expected for healthy animals (Figure 4A and B). After 72 h of UPEC infection, these compartments lose their punctate definition preferentially in the DNase II-deficient animals (Figure 4C and D). As the severity of this effect can be variable after infection, two animals are shown as representative examples for the AO staining with UPEC infection.

**Figure 4.**
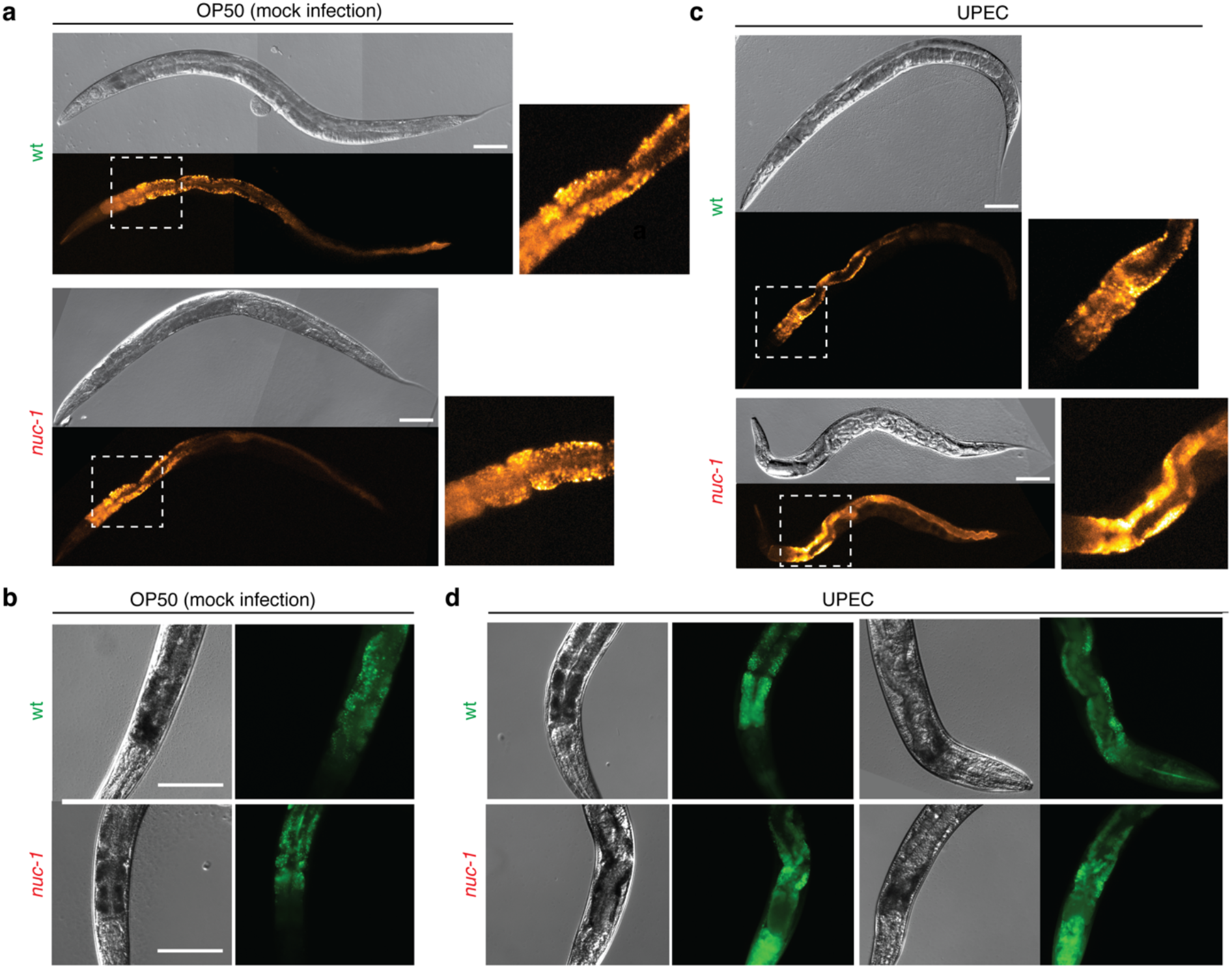
UPEC infection induces a loss of structural integrity in the intestine of *nuc-1* worms. **a.** Nile red staining of age-matched wt and *nuc-1* worms 72 hours after mock infection with OP50 or infection with UPEC at the L4 stage. **b.** Acridine orange staining of age-matched wt and *nuc-1* worms 96 h after mock infection with OP50 or infection with UPEC at the L4 stage. Images highlight the anterior (head) portion of the intestine. For all images, the scale bars represent 100 µm.

We used two approaches to examine body wall muscle integrity: locomotion (crawling speed) and phalloidin staining of F-actin (Francis, 1985; Wulf et al., 1979). As shown in Figure 5A, UPEC infection led to a more severe impairment of crawling speed in the DNase II-deficient worms compared to the wildtype worms. This effect could be due either to a behavioral change or to muscle degeneration. To distinguish these possibilities, we stained the F-actin in the body wall muscles used for crawling. Consistent with a decline in tissue integrity, the phalloidin staining revealed increased disorganization of the F-actin in the UPEC-infected worms compared to normally fed wildtype or DNase II-defective animals (Figure 5B). The white dashed lines outline individual muscle cells clearly showing atrophy and loss of F-actin organization most severely in the UPEC-infected, DNase II-defective animals.

**Figure 5.**
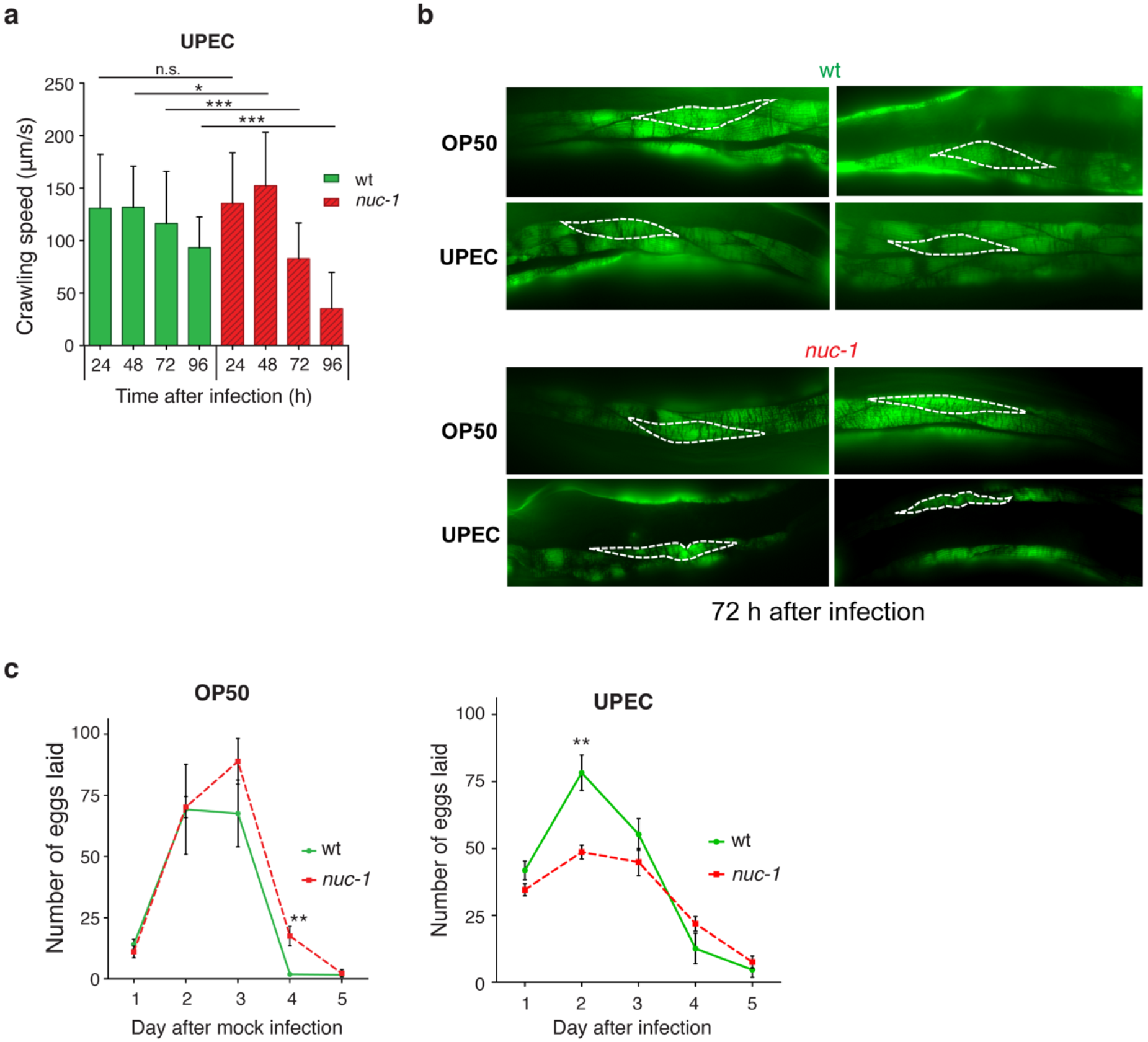
UPEC infection causes muscle degeneration and reproductive defects in DNase II-defective animals. **a.** Crawling speeds of UPEC-infected wt and *nuc-1* worms. **b.** Representative images of phalloidin-stained body wall muscles in mock-or UPEC-infected wt and *nuc-1* worms. White dashed outlines show individual muscle cells. **c.** Fecundity, scored as the number of eggs laid during the reproductive period, of wt and *nuc-1* worms mock-infected with OP50 or infected with UPEC. Shown are the means ± S.D. and statistical significance was assessed using unpaired, two-tailed Student’s *t* tests; ** 0.01 > P > 0.001.

Lastly, we examined reproductive function by measuring the fecundity of infected worms. While wildtype and DNase II-deficient worms have nearly identical progeny production when reared on *E. coli* OP50 (Figure 5C – left), UPEC-infected DNase II-deficient worms showed decreased egg production, especially during the first two days of egg laying (Figure 5C – right). Taken together, these results show that UPEC-infected DNase II-deficient worms suffer from profound systemic declines in tissue integrity and stability that is largely absent in the wildtype worms, suggesting that DNase II provides a protective effect during UPEC infection. Importantly, these phenotypes are reminiscent of the types of tissue decline associated with chronic inflammation in mammals indicating that *C. elegans* might be useful as a model for studying the effects of chronic inflammation associated with ectopic DNA.

### Disruption of the FSHR-1 innate immune signaling pathway rescues the tissue degeneration phenotype

One possibility to explain the observed tissue degeneration phenotypes is the aberrant induction of an uncontrolled immune response; if this hypothesis is true, then abrogating the response should alleviate the tissue decline in worms challenged with UPEC infection or cytoplasmic foreign DNA. The *C. elegans* p38 MAP kinase homolog PMK-1 is an important mediator of the innate immune response to a variety of pathogens (Kamaladevi and Balamurugan, 2015; Kim et al., 2002); thus, we first tested if it is involved in these phenotypes. RNAi-mediated depletion of PMK-1 did not rescue the pumping rate decline observed in *nuc-1* worms injected with CpG ODNs (Figure 6A). Thus, we conclude that p38/PMK-1 does not control the response to foreign cytoplasmic DNA.

**Figure 6.**
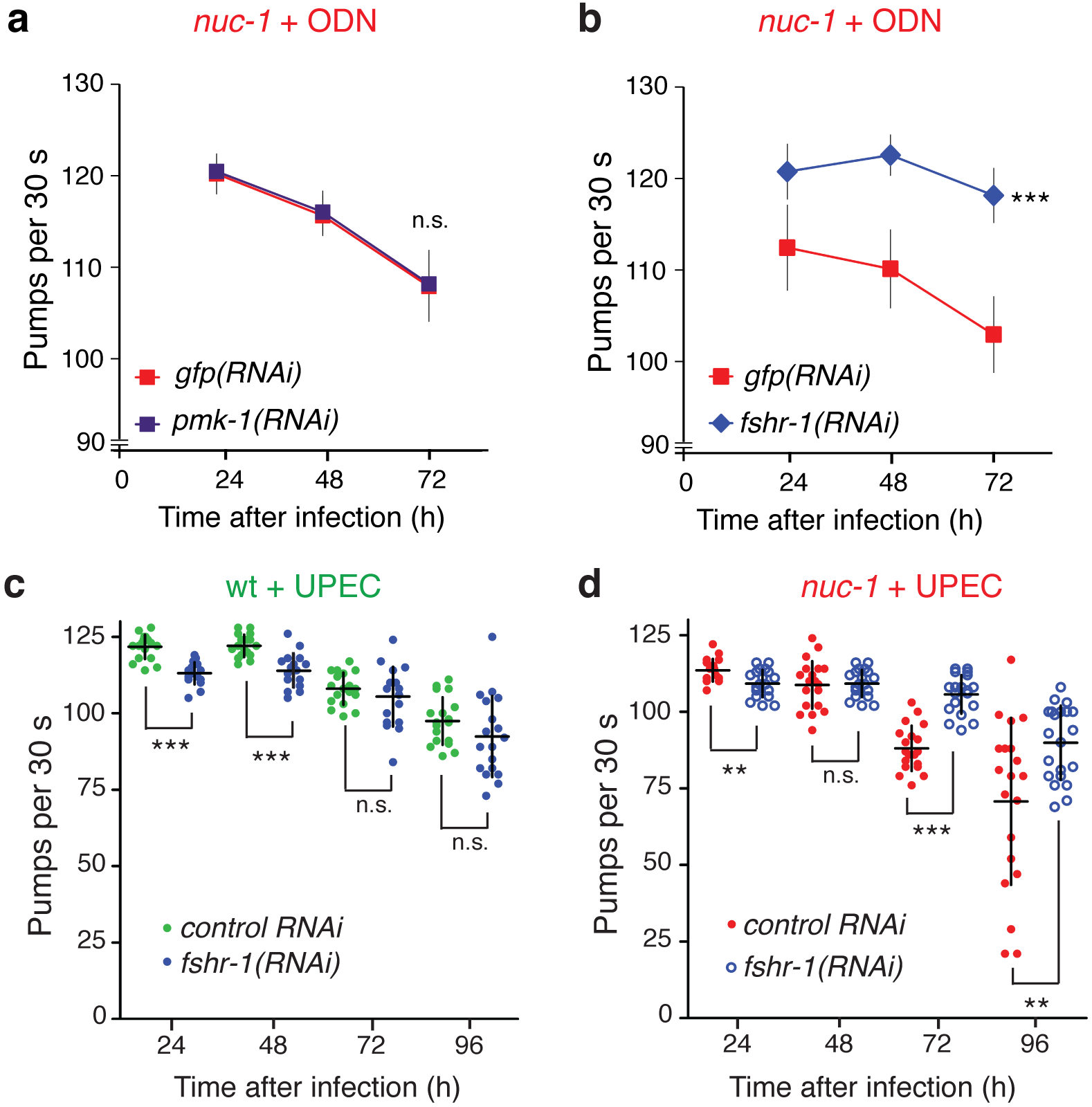
Tissue degeneration due to cytoplasmic DNA depends on immune signaling. **a. and b.** Plots of the pharyngeal pumping rates of *nuc-1* worms injected CpG-containing ODN with RNAi against *pmk-1* or *fshr-1.* **c. and d.** Plots of the pharyngeal pumping rates of wt and *nuc-1* worms infected with UPEC with and without RNAi against *fshr-1.* In all cases, the error bars show the means ± S.D. For a. and b., the statistical significance of differences in the trends between the control and experimental groups were assessed using two-way ANOVA. For c. and d., the statistical significance was assessed using unpaired, two-tailed Student’s *t* tests. In all cases, n.s. P > 0.05, * P < 0.01, ** 0.01 > P > 0.001, *** P < 0.001.

We next shifted our attention to another previously identified central immune regulator: the leucine-rich repeat (LRR)-containing protein FSHR-1, which regulates an immune response to several different pathogens in parallel to the p38/PMK-1 pathway (Powell et al., 2009). We tested for a role of FSHR-1 in our experimental system via RNAi-mediated depletion of FSHR-1 in wildtype and *nuc-1* worms. The worms were then either injected with CpG ODNs or infected with UPEC and the pharyngeal pumping rates were scored up to 96 h post-treatment. As shown in Figure 6B, FSHR-1 depletion in *nuc-1* worms eliminated the pumping rate decrease caused by injection of CpG ODNs into intestinal cells (Figure 6B). We also confirmed these phenotypes using UPEC infection. Depletion of FSHR-1 imparted no benefit to the UPEC-infected wildtype worms between 72 and 96 h (Figure 6C); in fact, the wildtype; *fshr-1(RNAi)* worms had decreased pumping rates at the early time points, suggesting that FSHR-1 might control a rapid beneficial response to the pathogen. In contrast, UPEC-infected *nuc-1; fshr-1(RNAi)* worms showed robust improvements in their pumping rates at the later time points of infection that were similar to wild-type rates (Figure 6D). As in the wildtype worms, FSHR-1 depletion in *nuc-1* worms also had a slight negative effect on pumping rate at the 24 h time point. To further confirm the presence of an innate immune response, we tested the expression levels of several previously identified immune-associated genes by RT-qPCR and found that their expression was indeed induced, both after UPEC infection and DNA injection (Supplemental Figure 2). Taken together, these results suggest that FSHR-1 regulates a beneficial early response to infection that is common between wildtype and *nuc-1* worms (most likely independent of foreign DNA), but that FSHR-1 activity is the primary, if not the sole, mediator of the response to persistent foreign cytoplasmic DNA.

### Chronic UPEC infection leads to a disruption of protein homeostasis

We next set out to clarify the basis of the increased sensitivity of the DNase II-deficient worms to cytoplasmic DNA (either directly injected or introduced via UPEC infection). Immunity-associated stress responses in *C. elegans* result in induction of a large number of putative secreted peptides (Shivers et al., 2008) and deregulation of the response can be detrimental to *C. elegans* (Cheesman et al., 2016). One hypothesis to explain the preferential tissue degeneration in DNase II-deficient worms is that persistent cytoplasmic DNA leads to a prolonged upshift in the production of secreted proteins that could strain the animal’s protein quality control mechanisms to the point that proteotoxic effects cause cellular damage. This hypothesis is supported by the previous observation that the UPR^ER^ is required by worms to tolerate the innate immune response to the pathogen *Pseudomonas aeruginosa* (Richardson et al., 2010), and that the UPR may assist in activation of the immune response (Dai et al., 2015).

As our first approach to look for effects on protein homeostasis, we used transmission electron microscopy to directly examine the ER structure and ribosome content in the intestinal cells. As can be seen in Figure 7A, the ER of wildtype worms, either mock infected with *E. coli* OP50 or infected with UPEC, appears normal with a regular distribution of ribosomes around the ER cisternae (see high magnification images, blue arrows). In contrast, the DNase II-deficient worms infected with UPEC showed enlarged cisternal spaces, a typical hallmark of ER disruption (Richardson et al., 2010). Furthermore, the distribution of the ribosomes around the cisternae was irregular (compare regions indicated by red arrows to those indicated by blue arrows), especially in infected DNase II-deficient worms.

**Figure 7.**
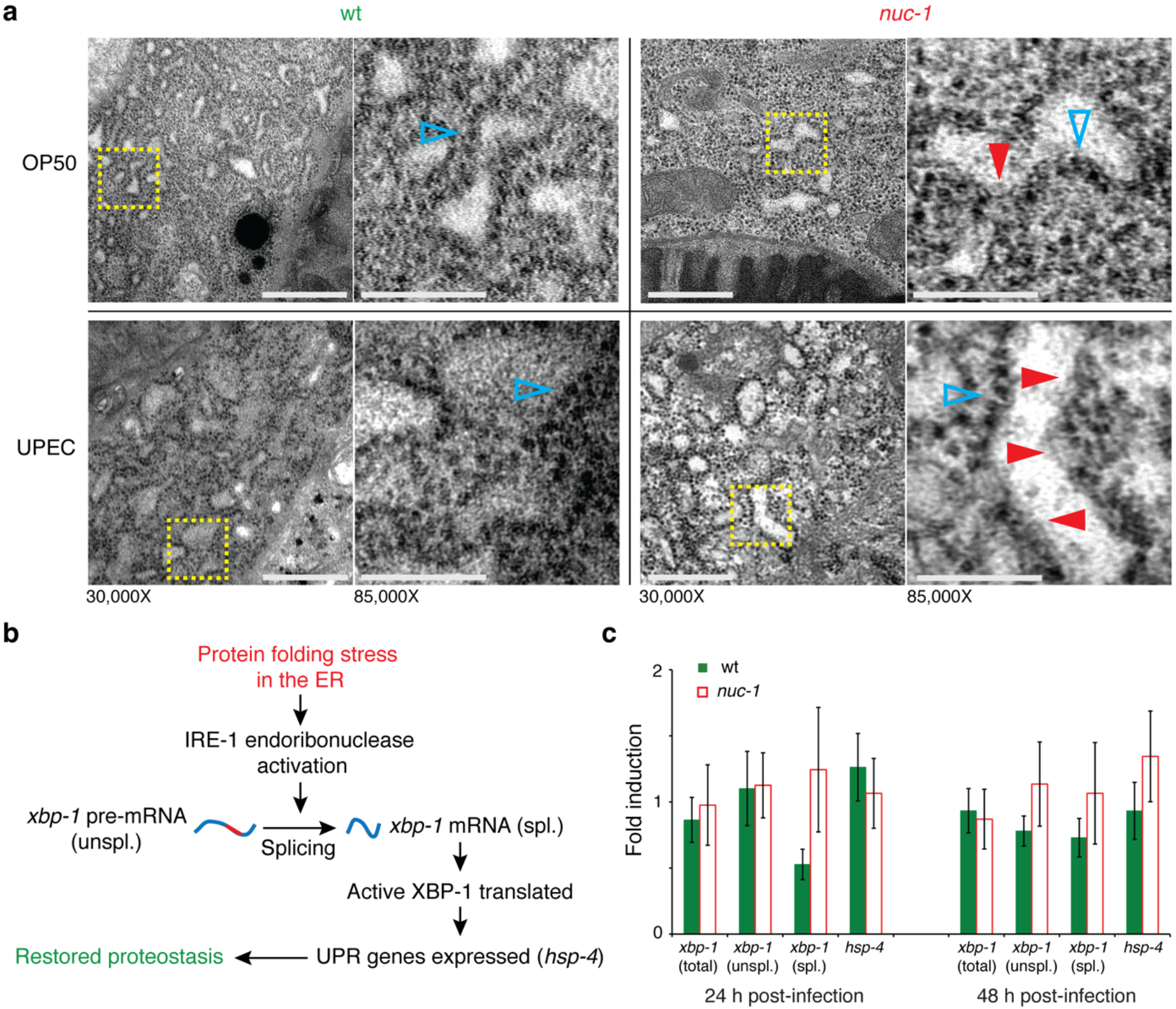
UPEC infection leads to endoplasmic reticulum disruption in *nuc-1* worms with no compensatory induction of the UPR^ER^. **a.** Transmission electron micrographs showing the endoplasmic reticulum in mock-infected and UPEC-infected wt (left) and *nuc-1* (right) worms. Magnifications are indicated and the scale bars represent 500 nm. Yellow boxes highlight areas shown in higher magnification. Open blue arrows indicate regions of the ER with normal ribosome distribution; filled red arrows indicate regions with disrupted ribosome distribution. **b.** Schematic of the UPR^ER^ pathway in *C. elegans.* Abbreviations correspond to those in c. **c.** Plot showing the relative levels of different forms of the *xbp-1* transcript 24 and 48 h post-infection: *xpb-1* total, *xbp-1 xbp-1* pre-mRNA (unspl.), *xbp-1* spliced (spl.), and the XBP-1 regulatory target *hsp-4.* Data shown are the means ± S.D. No statistically significant differences were observed.

Physical disruption of the ER could be associated with defects in protein folding, which would normally lead to compensatory activation of the UPR^ER^. To evaluate the status of the UPR^ER^, we measured the relative levels of the unspliced (unspl.) and spliced (spl.) forms of the *xbp-1* transcript (Yoshida et al., 2001). Splicing of *xbp-1* leads to the production of the active form of XBP-1, a transcription factor that acts as a central regulator of the UPR^ER^ (Figure 7B). RNA was collected from wildtype and DNase II-deficient worms 24 and 48 h after infection with UPEC. As shown in Figure 7C, the level of spliced *xbp-1* transcript is not significantly increased under any condition and, importantly, the expression of *hsp-4,* a prototypical XBP-1 target gene (see Figure 7B), is also not induced. These results suggest two mutually exclusive possibilities: (1) Despite disruption of the ER, no compensatory activation of the UPR^ER^ occurs, or (2) that the response is activated, but is not sufficiently sustained during the course of the infection to provide a long-term protective effect. We conclude from these results that a response by the DNase II-deficient worms to UPEC infection strains the protein quality control pathways leading to unresolved protein-folding stress in the ER (due to failure to activate the UPR^ER^) and that ensuing proteotoxic effects could ultimately lead to the tissue degeneration in DNase II-deficient animals due to the accumulation of cytoplasmic DNA. These results are consistent with the toxicity of an aberrantly activated innate immune response in *C. elegans* (Cheesman et al., 2016).

### Therapeutic improvement of ER protein homeostasis alleviates the tissue functionality decline caused by persistent cytoplasmic DNA

If a chronic, unresolved defect in ER homeostasis due to failure to induce or sustain the UPR^ER^ drives the tissue degeneration in the presence of persistent cytoplasmic DNA, we wondered if exogenous, sustained activation of the UPR^ER^ or enhancement of the ER protein folding capacity could alleviate the tissue degeneration. To examine these possibilities, we used three distinct approaches: (1) ectopic activation of the UPR^ER^ via mild, subtoxic, chemically-induced protein folding stress (Figure 8A); (2) activation of the UPR^ER^ independent of protein folding stress (Figure 8D); and (3) therapeutic enhancement of protein folding capacity in the ER (Figure 8F).

**Figure 8.**
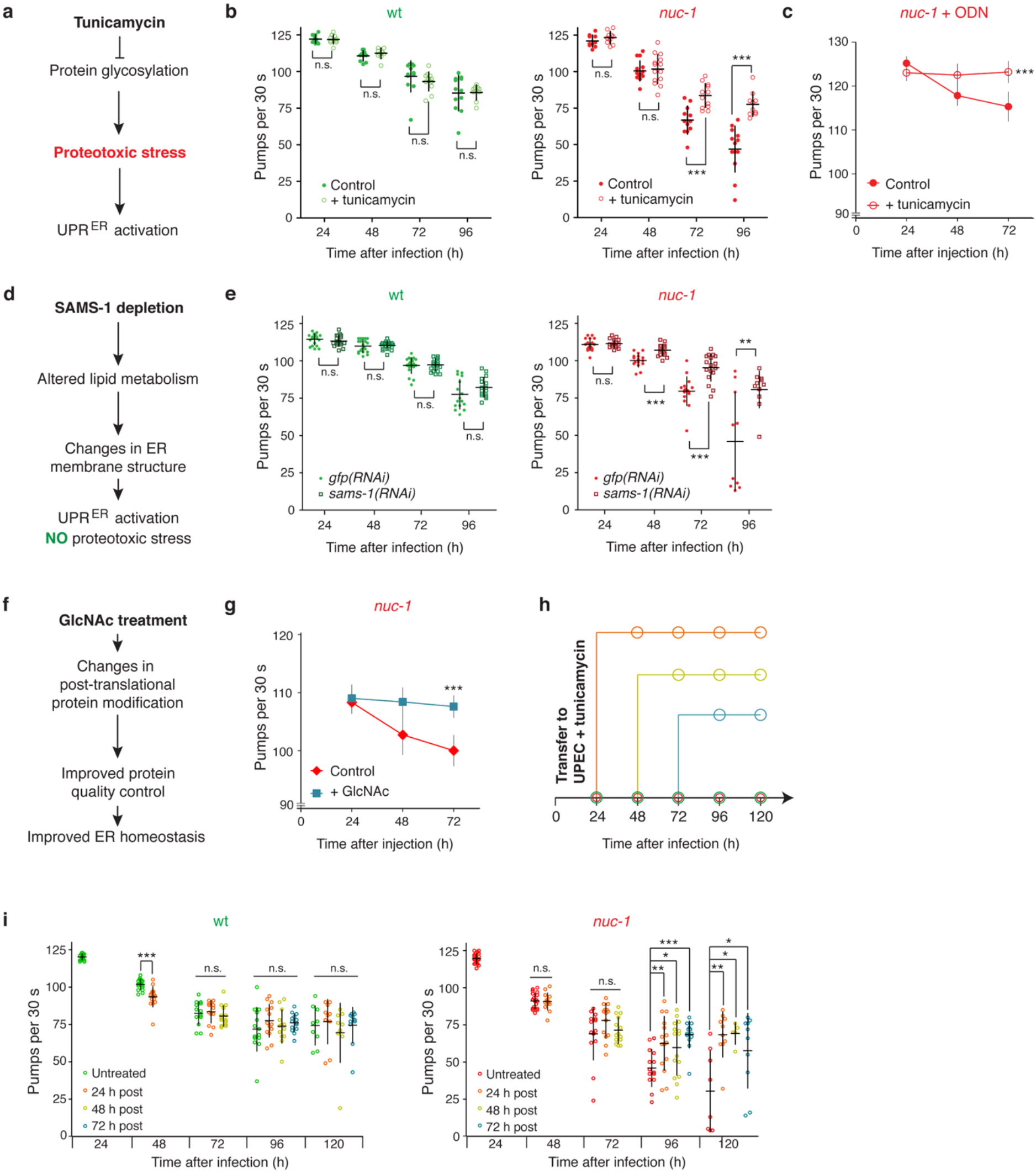
Elevated protein homeostasis can alleviate tissue functionality declines due to UPEC infection and ODN injection. **a.** Schematic of UPR^ER^ activation by tunicamycin. **b.** Plots of the pharyngeal pumping rates of wt and *nuc-1* worms infected with UPEC with and without treatment with 0.05 μg/mL tunicamycin. **c**, Plot of the pharyngeal pumping rates of worms injected with ODNs with and without treatment with 0.05 μg/mL tunicamycin. **d.** Schematic of UPR^ER^ activation by *sams-1* depletion. **e.** Plots of the pharyngeal pumping rates of wt and *nuc-1* worms infected with UPEC after treatment with control RNAi *(gfp)* or RNAi against *sams-1.* **f.** Schematic of ER stabilization by GlcNAc. **g.** Plot of the pharyngeal pumping rates of worms injected with ODNs with and without treatment with 10 mM GlcNAc. **h.** Outline of the experiment shown in i, where the colors correspond to data points in the following plots. **e**, Plots of the pharyngeal pumping rates of wt and *nuc-1* worms infected with UPEC without treatment and with treatment with 0.05 μg/mL tunicamycin at the time points shown in h. All plots show the means ± S.D. For **b**, **e**, and **i**, statistical significance was assessed using unpaired, two-tailed Student’s *t* tests; n.s. P > 0.05, * P < 0.01, ** 0.01 > P > 0.001, *** P < 0.001. For **c** and **g**, the statistical significance of the effect was assessed using two-way ANOVA (*** P < 0.001).

We first used a continuous low dose (0.5 μg/mL) treatment with tunicamycin during the course of UPEC infection and after DNA injection. Tunicamycin inhibits protein glycosylation, which leads to subsequent induction of the UPR^ER^ (King and Tabiowo, 1981)(Figure 8A). Wildtype worms treated with tunicamycin during UPEC infection showed no improvement in pharyngeal pumping over 96 h of infection, although they still showed an infection-dependent health decline (Figure 8B – left). In stark contrast, tunicamycin treatment of UPEC-infected DNase II-deficient worms reverted the decreased pumping rate to a level similar to infected wildtype worms (Figure 8B – right). Remarkably, tunicamycin treatment also completely eliminated the pumping rate decline caused by ODN injection into the intestine of DNase II-deficient animals (Figure 8C). These results, in particular the rescue of the injection effect, suggest that exogenous activation of the UPR^ER^ can resolve the detrimental consequences of persistent cytoplasmic DNA.

Because tunicamycin interferes with post-translational processing and can be toxic (Denzel et al., 2014; King and Tabiowo, 1981), we wanted to confirm that its beneficial effect was due the induction of UPR^ER^ and not to off-target effects. To this end we sought an alternative experimental strategy to activate the UPR^ER^ without affecting the protein homeostasis systems. Loss of the S-adenosylmethionine synthase SAMS-1 leads to induction of the UPR^ER^ by inducing lipid disequilibrium in the ER without introducing protein folding stress (Hou et al., 2014) (Figure 8D). Mirroring the tunicamycin results, RNAi-mediated depletion of *sams-1* led to a similar rescue of the decline in tissue functionality after UPEC infection in DNase II-deficient worms and provided little or no benefit for wildtype worms (Figure 8E). While *sams-1* has been previously assigned a function in immunity (Ding et al., 2015), a general change in proteostatic status is unlikely here since the effect of infection on the wildtype strain was minimal, and the beneficial effect was only observed in the DNase II-deficient strain. We conclude from these experiments that mild ectopic induction of the UPR^ER^ can relieve the proteotoxic outcomes from an upshift in protein synthesis associated with the worm’s stress response pathways.

Tunicamycin treatment and SAMS-1 depletion activate the UPR^ER^ via disruption of normal cellular processes (protein glycosylation and lipid homeostasis, respectively). We next wondered if protection of ER function without such side effects could also provide protection against tissue degeneration in DNase II-deficient worms. Treatment of *C. elegans* with *N*-acetylglucosamine (GlcNAc) slows ageing and suppresses proteotoxic effects of aggregating proteins by augmenting the hexosamine pathway, which enhances protein folding and improves ER homeostasis (Denzel et al., 2014) (Figure 8F). To test whether GlcNAc treatment can protect DNase II-deficient worms from the detrimental effects of cytoplasmic DNA, we treated DNase II-deficient animals with 10 mM GlcNAc after ODN injection. As shown in Figure 8G, constant exposure to GlcNAc following the ODN injection eliminated the tissue functionality decline caused by DNA injection; thus, enhancement of the ER protein folding capacity through this non-toxic pharmacological approach can offer a similar benefit to tunicamycin treatment and SAMS-1 depletion.

Finally, we were interested in whether tunicamycin treatment could be used as a curative intervention to provide a therapeutic benefit during ongoing infection. To test this possibility, we transferred infected worms from plates containing only UPEC to plates that additionally contain 0.5 μg/mL tunicamycin at regular intervals up to 72 h post-infection and measured the pumping rates on the following days up to 120 h post-infection (outlined in Figure 8H). Remarkably, DNase II-deficient worms transferred to tunicamycin plates as late as 72 h post-infection showed significant improvement in the pumping rates (Figure 8I – right). Further supporting the conclusion that wildtype worms are not subject to the same unresolved ER stress, transfer to tunicamycin offered them no advantage (Figure 6I – left). Furthermore, this increase in health was visible by gross examination of the worms. As shown in Supplemental Figure 3B, DNase II-deficient worms transferred to tunicamycin 72 h post-infection and examined 120 h post-infection (the latest time of treatment and endpoint of the experiment) showed signs of physical health absent in the untreated worms, most notably the typical sinusoidal waveform for the body present during normal crawling. No obvious differences in the physical attributes of the wildtype worms were visible following treatment, as they are, in general, resistant to the UPEC infection (Supplemental Figure 3A). Based on these results, we conclude that mild induction of the UPR^ER^ can provide a protective effect during an ongoing immunogenic challenge that induces proteotoxic stress.

## DISCUSSION

Many open questions remain about the mechanisms of sensing and responding to cytoplasmic pathogenic DNA. Perhaps more enigmatic are the mechanisms through which chronic innate immune responses lead to tissue damage. Furthermore, our repertoire of therapeutic options for alleviating these detrimental outcomes is limited. In this study, our goal was to establish *C. elegans* as a simple metazoan model to further explore how animals respond to cytoplasmic DNA and to better understand the effects chronic activation of the downstream responses at the cellular and organismal levels. Our ultimate goal was to develop interventions to support tissue preservation and functionality under such conditions.

The nematode innate immune response has been extensively investigated, and several key regulators that mediate systemic responses to bacteria and fungi have been well documented (Ermolaeva and Schumacher, 2014; Kim et al., 2002; Kim and Ewbank, 2018; Powell et al., 2009). However, what is entirely lacking in the *C. elegans* field are data on whether the worm can sense and respond to cytoplasmic DNA. None of the known DNA-sensing pathways identified in higher eukaryotes are conserved in *C. elegans.* While the *C. elegans* genome does encode one Toll-like receptor (TOL-1) (Pujol et al., 2001), it appears to have a limited role in immunity (Battisti et al., 2017; Galbadage et al., 2016; Rangan et al., 2016; Tenor and Aballay, 2008); furthermore, the immune functions it does have may operate through a developmental function (Brandt and Ringstad, 2015). The absence of these systems suggests that *C. elegans* may possess alternative and novel pathways to fulfill this important defense function. In this study, we demonstrated that *C. elegans* can indeed mount a response to foreign cytoplasmic DNA, and that when this response is chronically activated it can lead to severe multi-system declines in tissue stability and functionality. After identifying the sub-cellular effects responsible for these declines, we successfully developed simple interventions that alleviate these effects, which may also be applicable to higher organisms, including humans. It is this novel therapeutic perspective supplied by our study that we believe could have the most impact on human health as these approaches are refined.

Upon introduction of foreign DNA into the cytoplasm of worm intestinal cells, either by direct microinjection or via infection with a uropathogenic *E. coli* (UPEC) strain isolated from a human patient (Figures 1, 2 and 3), we observed only a mild effect after UPEC infection and almost no effect after DNA injection in wild type worms. The decreased survival caused by the UPEC strain is likely due to effects of bacterial colonization of the intestine and incomplete host-mediated immunity. However, a different story emerged in a strain lacking DNase II, encoded by *nuc-1* in *C. elegans.* Introduction of pathogenic DNA into the cytoplasm of the intestinal cells these worms led to profound declines in tissue stability and functionality that worsened over the course of several days. An unexpected outcome of the introduction of foreign DNA was that tissues distal to the intestinal, including the pharynx (Figures 1 and 3), the muscles (Figure 5A and B), and the reproductive system (Figure 5C), were severely affected. One explanation for these phenotypes is that the expression levels of many secreted factors increase during the nematode innate immune response (Kato et al., 2002; Mallo et al., 2002; Schulenburg et al., 2004) and that these factors can, when chronically expressed, have detrimental effects throughout the body. We propose that these effects are reminiscent of the tissue damage that occurs during chronic inflammatory responses in mammals, including humans.

Dysregulation of the p38/PMK-1-mediated response to *P. aeruginosa,* in the form of chronic induction, is toxic to worms (Cheesman et al., 2016). Moreover, in developing worms the UPR^ER^ is required for survival during *P. aeruginosa* infection (Richardson et al., 2010). These observations prompted us to investigate whether the chronic response to pathogenic DNA could disrupt protein homeostasis. To explore this question, we used transmission electron microscopy to examine the structure of the ER (Figure 7A) and molecular and cellular techniques to examine the function of the UPR^ER^ (Figure 7B). We observed that not only was the superstructure of the ER disrupted (a previously characterized feature of impaired protein homeostasis (Richardson et al., 2010), but that the UPR^ER^ was not active during chronic (on the scale of days) exposure to cytoplasmic DNA. One possible explanation that we favor is that the UPR^ER^ may be activated during the initial phase of the response, but that it cannot be maintained when the immune response becomes chronic. The lack of this ongoing compensatory response as well as the burden of the enhanced gene expression likely lead to proteotoxic effects that we believe are the sub-cellular basis for the observed tissue degeneration.

We also explored interventions that if successful in alleviating these detrimental effects, would substantiate the functional role of the UPR^ER^ and potentially precipitate novel anti-inflammatory therapeutic routes. First, we attempted to entirely ablate the underlying immune response by eliminating the most likely immune regulators (Figure 6). We depleted two of the key response regulators, PMK-1 and FSHR-1, and challenged the worms with pathogenic cytoplasmic DNA. Loss of PMK-1 in the DNase II-deficient worms had no effect on the tissue degeneration phenotype; in contrast, loss of the G protein-coupled receptor FSHR-1 (Powell et al., 2009) robustly alleviated the phenotype. Through this experiment, we confirmed that chronic activation of the FSHR-1-mediated immune response indeed underlies the tissue degeneration phenotype, and we propose FSHR-1 as the likely mediator in the response to pathogenic cytoplasmic DNA. Given the structure of the leucine-rich repeats in FSHR-1 that are also present in mammalian toll-like receptors, it is not inconceivable that this immune signaling component might even function as sensor of cytoplasmic DNA. Based on our observation of ER disruption and a lack of sustained UPR^ER^ induction (Figure 7), we used three mechanistically independent approaches that improve protein homeostasis with the goal of reducing the toxicity of the chronic response (Figure 8). We exposed the worms to low, sub-toxic levels of tunicamycin, a chemical that robustly activates the UPR^ER^ by interfering with *N*-linked glycosylation of newly translated proteins. This treatment induced a profound rescue of the tissue degeneration phenotype in the DNase II-deficient worms exposed to pathogenic DNA. We suggest that this treatment constitutively activates and maintains the UPR^ER^ during the entire course of the chronic response, and that this ongoing compensatory response rescues the tissue degeneration phenotype. As an alternative genetic approach, we depleted the S-adenosyl methionine synthetase SAMS-1 via RNA interference. Loss of SAMS-1 is known to enhance protein homeostasis without inducing proteotoxic stress (in contrast to tunicamycin). Similarly, SAMS-1 depletion also rescued the tissue degeneration phenotype. As both tunicamycin exposure and SAMS-1 depletion inhibit fundamental cellular processes to induce the UPR^ER^, we wanted to explore a completely non-toxic approach. Treatment with *N*-acetylglucosamine (GlcNAc) enhances protein homeostasis by increasing the protein folding capacity of the ER. Similar to the other two interventions, GlcNAc treatment also rescued to the time-dependent tissue degeneration phenotype. Finally, in addition to the studies by Richardson *et al.* and Dai *et al.* cited above, protein homeostasis has been further identified as important component in the response to infection in *C. elegans* (Richardson et al., 2010). Furthermore, enhanced ER homeostasis has recently been shown to enhance the animals’ tolerance of the innate immune response (Tillman et al., 2018). These outside observations further substantiate our model, which is outlined in Figure 9.

**Figure 9.**
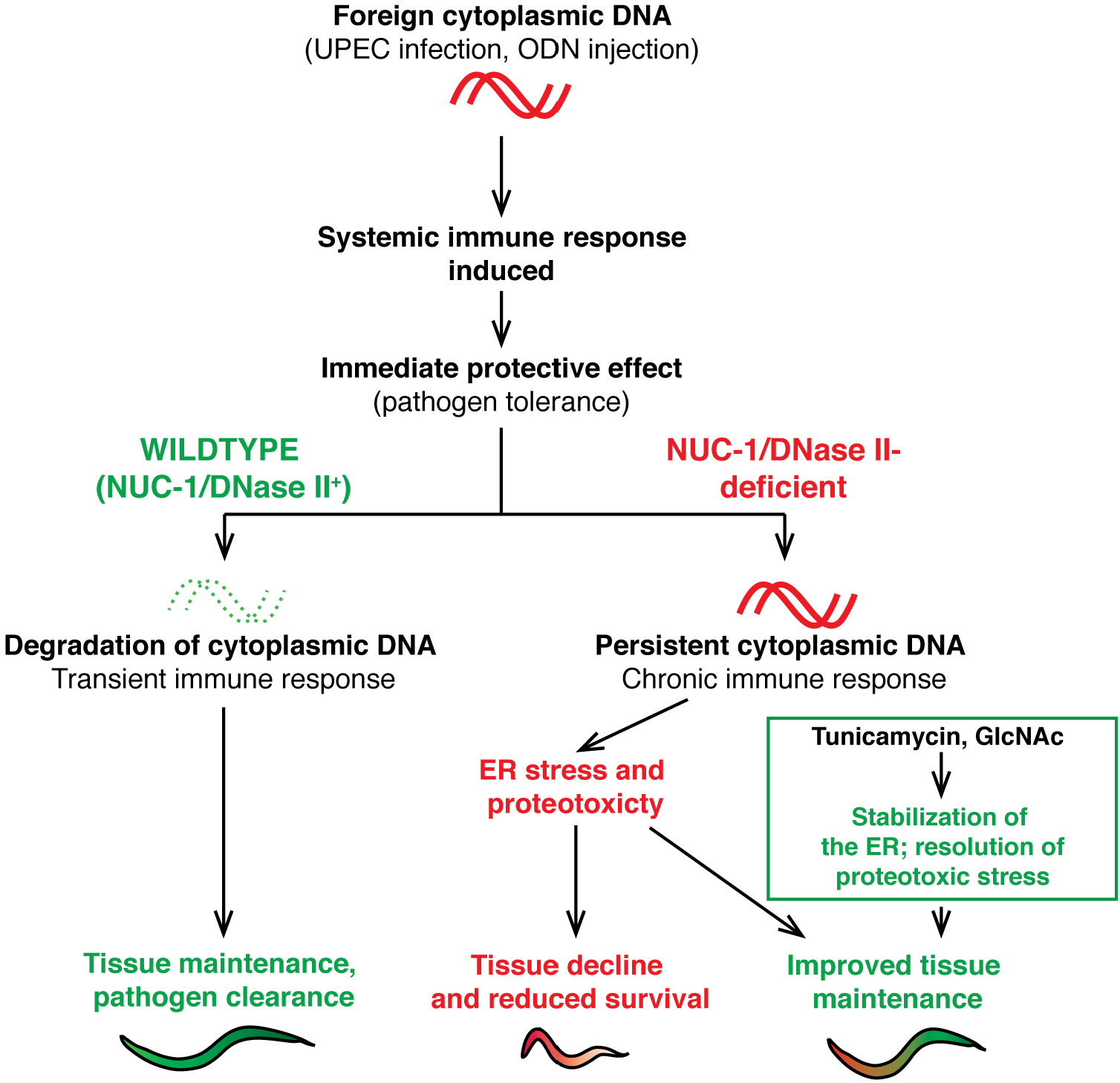
Model for the tissue functionality declines caused by persistent cytoplasmic DNA. Cytoplasmic DNA triggers an immune response via FSHR-1 that provides early protection against infection; however, when the foreign DNA persists as a result of DNase II-deficiency, the immune response becomes chronic and results in an inflammation-like response that leads to disruption of the ER leading to declines in tissue functionality and ultimately reduced survival. Therapeutic activation of the endoplasmic reticulum unfolded protein response or augmentation of protein folding by GlcNAc can rescue these tissue declines.

We believe that the identification of therapeutic approaches that alleviate the outcomes of the chronically activated innate immune response may have the most importance from the perspective of human health. Treatment with low concentration of tunicamycin (which has been shown to be well tolerated by mice (Wang et al., 2015)) or GlcNAc had robust protective effects in the DNaseII-deficient worms following UPEC infection or intestinal DNA injection. The dependence of this rescue on the absence of DNase II together with our ability to rescue the phenotype via disruption of innate immune signaling, strongly supports that these outcomes are, indeed, a direct result of a systemic response to the pathogenic DNA; thus, we demonstrate for the first time that *C. elegans* does mount a response to cytoplasmic DNA. Furthermore, we propose that the tunicamycin-induced protection occurs via low-level UPR^ER^ activation, and that the enhanced protein folding capacity afforded by GlcNAc helps to maintain ER functionality. The net outcome of each interventions is an amelioration of the proteotoxic effects resulting from the chronic immune response. These findings may have important implications for the understanding of *DNASE2*- related inflammatory conditions in humans. ER stress has been shown to be a mediator of kidney injury (reviewed in (Inagi, 2009)), and tunicamycin injection reduced the severity of the inflammatory kidney disease in a mouse model for mesangioproliferative glomerulonepthritis (Inagi et al., 2008). These observations provide additional support for our hypothesis that mild UPR^ER^ induction may provide protection in the face of the significant upshift in protein expression involved in inflammatory responses. Therefore, we propose that therapeutic protection of ER functionality in patients with inflammatory conditions, such as rheumatoid arthritis and inflammation-induced heart failure, may offer a similar benefit.

The innate immune response also puts significant strain on the host’s cells and the ensuing inflammation can result in tissue damage (Wallach et al., 2014). Inflammation can disrupt tissue functionality and, when chronically active, can contribute to the aging process (Pawelec et al., 2014). Inflammatory processes are associated with a wide range of human disorders including bowel disease (Hanauer, 2006), arthritis (Firestein, 2003), and heart failure (Oka et al., 2012). It is, thus, of pivotal importance to better understand the molecular mechanisms how the innate immune responses result in inflammation. The characterization of this new tissue degeneration phenotype in worms, which we propose can serve as a model for inflammation-associated tissue degeneration in higher animals, as well as our identification of a previously unknown response to persistent cytoplasmic pathogenic DNA establishes *C. elegans* as an ancestral model for investigating inflammatory responses and for determining the role of protein quality mechanisms in counteracting the detrimental consequences of inflammation. We suggest that further exploration of these processes in this metazoan model will provide new perspectives into the cellular and organismal responses to pathogenic DNA and persistent challenges to the innate immune system.

## ACKNOWLEDGEMENTS

We are grateful to Jennifer Engelmeyer for outstanding technical support in the laboratory and to Astrid Schauss and Beatrix Martiny (CECAD Imaging Facility) for sample preparation for the transmission electron microscopy. Worm strains were provided by the *Caenorhabditis* Genetics Center (funded by the NIH National Center for Research Resources) and the National Bioresource Project. The UPEC strain was provided by Olaf Utermöhlen (Institute for Medical Microbiology, Immunology and Hygiene, Medical Center, University of Cologne). BS acknowledges funding from the Deutsche Forschungsgemeinschaft (CECAD, SFB 829, SFB 670, KFO 286 and KFO 329), the European Research Council (ERC Starting grant 260383), the Deutsche Krebshilfe (70112899), the COST action (BM1408, GENiE), and the Bundesministerium für Forschung und Bildung (Sybacol FKZ0315893). The authors declare no competing interests.

## AUTHOR CONTRIBUTIONS

A.B.W. designed and led the project, performed experiments, and, with B.S., wrote the manuscript. J.- E.M. performed experiments and assisted with data analysis. F.H. performed the injection experiments and assisted with data analysis. W.B. performed the transmission electron microscopy and assisted with the image analysis. B.S. supervised the project and collaborated on writing the manuscript. All of the authors were involved in critically reviewing and editing the manuscript.

---

**Supplemental Figure 1.**
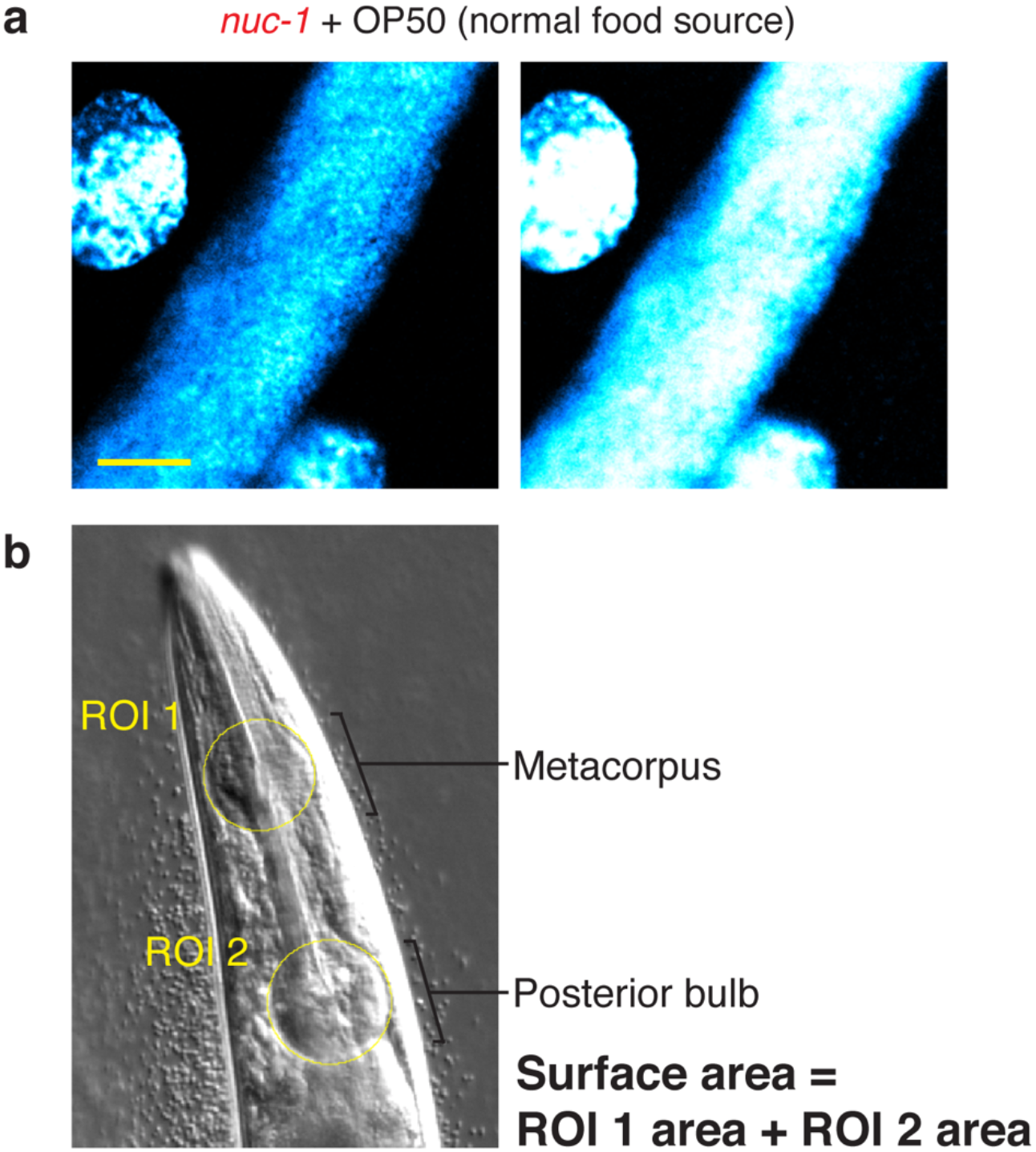
Controls for the cytoplasmic DNA imaging in Figure 2 and explanation of pharynx size measurement in Figure 3. **a.** DAPI staining of *nuc-1* worms mock-infected with OP50. **b.** Diagram of the method used to calculate the surface area of the pharyngeal bulbs (see *Experimental Procedures*).

**Supplemental Figure 2.**
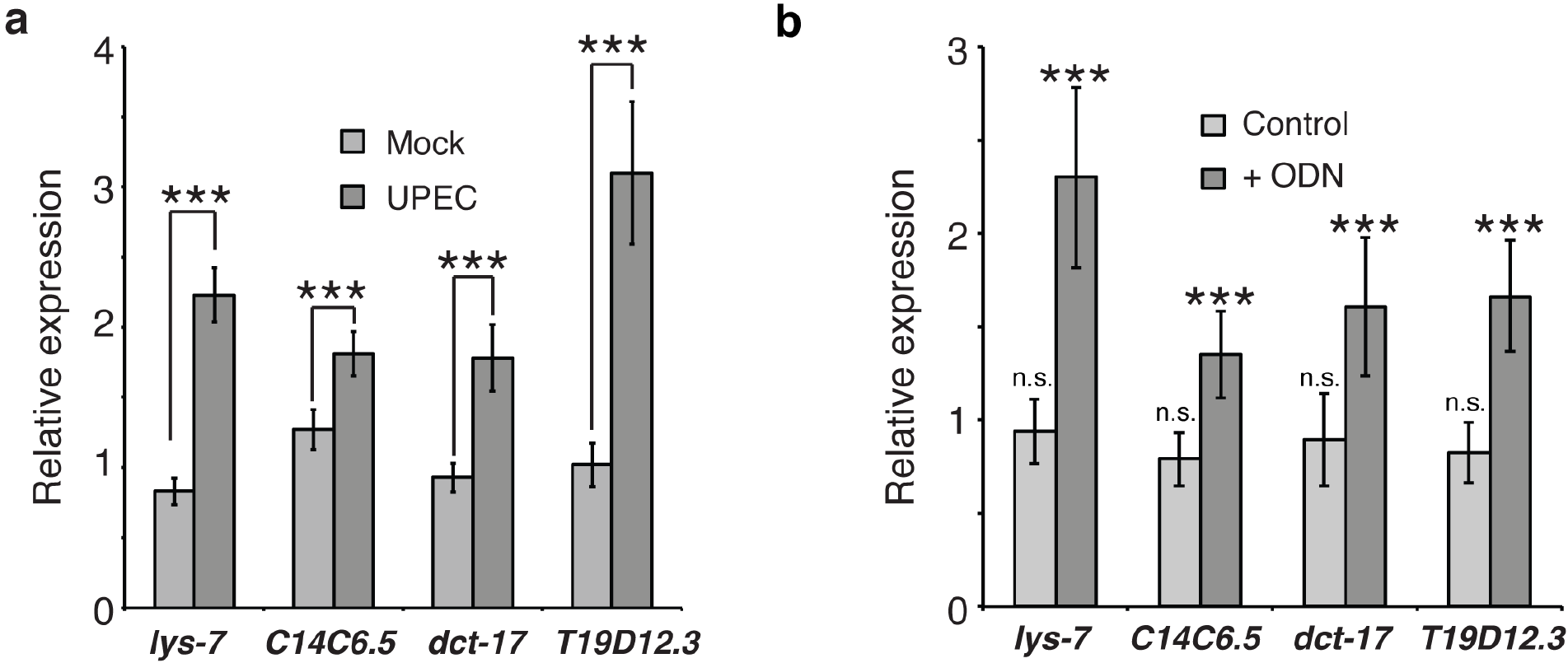
Supplemental data for the experiments in Figure 6. Cytoplasmic DNA preferentially induces immune gene expression in *nuc-1* worms. **a.** RT-qPCR analysis of the relative expression of immune genes comparing *nuc-1* to wt following mock-infection or UPEC infection. **b.** RT-qPCR analysis of the relative expression of immune genes following ODN injection comparing *nuc-1* to wt worms.

**Supplemental Figure 3.**
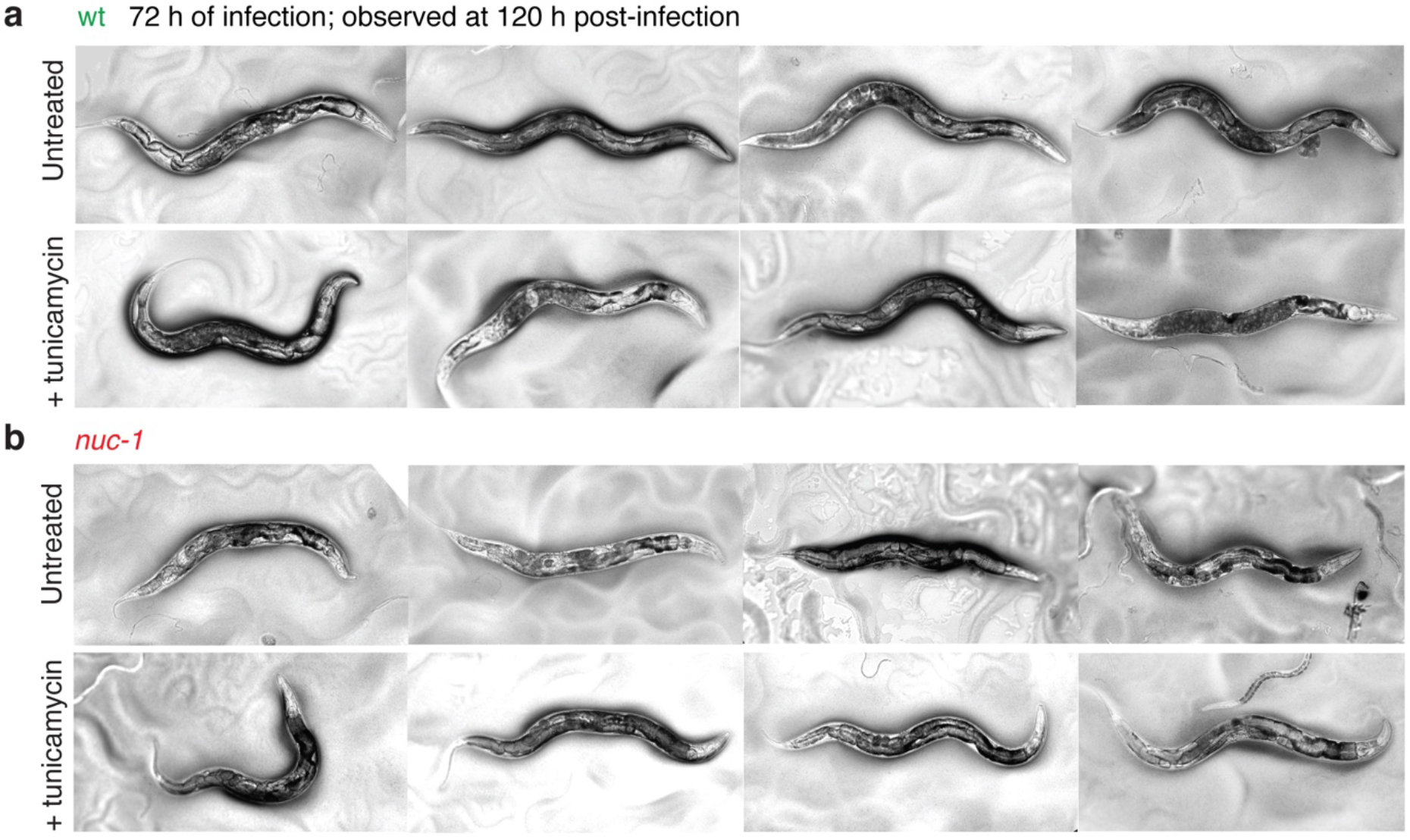
Supplemental data for therapeutic treatment experiments in Figure 8. Representative images of wt (**a**) and *nuc-1* (**b**) worms infected with UPEC for 72 h before 48 h of tunicamycin treatment (0.05 μg/mL) with continuous UPEC exposure.

## EXPERIMENTAL PROCEDURES

### Strain maintenance

Unless otherwise indicated (see pathogen infection methods), all strains were maintained as described at 20 °C (Brenner, 1974). *E. coli* strain OP50 was used for routine feeding. The N2 Bristol strain is used as the wildtype control. Other strains used in this study:

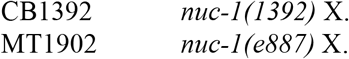

Except for where indicated in Figure 3, the *nuc-1(e1392)* allele is used throughout.

### RNAi treatment

RNAi feeding vectors for *fshr-1*, *pmk-1*, and *sams-1* were obtained from the Ahringer RNAi library (Source Bioscience, Nottingham, United Kingdom). RNAi feeding *E. coli* strains were cultured as follows: An overnight culture in LB liquid supplemented with ampicillin (0.1 mg/mL) was inoculated from a frozen stock and incubated for 37 °C overnight with vigorous aeration. This culture was diluted 1:1000 in fresh medium and grown to an OD_600_ of about 0.6 under similar conditions. At this time, IPTG was added to a final concentration of 2 mM and the cultures were incubated for an additional 3 h. 0.2 mL of this culture was then added to NGM plates supplemented with the same concentration of IPTG and grown overnight at 37 °C. For *sams-1,* L2/L3 worms were fed RNAi for 10 hours prior to infection as L4.

### Microinjection

Worms were picked the day before injection at L4 larval stage and grown overnight at 20°C on NGM plates seeded with OP50. The general methodology followed is described in detail in (Rieckher and Tavernarakis, 2017). The next morning, the young adult worms were transferred into halocarbon oil (Sigma, St. Louis, MO) on a 2% agarose pad for immobilization and injection. Microinjections were performed using an Axio Observer A1 (Zeiss) and FemtoJet (Eppendorf, Hamburg, Germany) microinjection system. Femtotips II (Eppendorf) were used for the injection and the fluorescent dye conjugate rhodamine B isothocyanate-dextran (10,000 MW) (Sigma) was used to track the injection at a concentration of 6 mg/mL in egg salts buffer. DNA from the pathogenic E. *coli* strain ECO85 was purified using the Puregene Core Kit A (Qiagen, Hilden, Germany) and injected at a concentration of 8 μg/μl and the synthetic CpG oligodeoxyribonucleotide ODN 2395 (Miltenyi Biotec, Bergisch Gladbach, Germany) was injected at a concentration of 100 ng/μl. All injections were made directly into a single intestinal cell. Following injection, the worms were washed with a mixture of M9 buffer and antibiotics (0.1 mg/mL streptomycin and 0.1 mg/mL gentamycin) to minimize contamination. 50 μl of the antibiotic solution was placed onto an empty NGM plate and up to 20 worms were placed in the liquid, which was allowed to completely soak into the agar. The worms were then transferred onto NGM plates seeded with *E. coli* OP50. Next, the worms were sorted using an Axio Zoom.V16 (Zeiss, Jena, Germany) and only worms containing rhodamine B isothiocyanate-dextran in intestinal cells (and not in the pseudocoelomic space or intestinal lumen) were kept for further experiments. Each selected worm was placed on an individual NGM plate seeded with OP50 and maintained at 20 °C.

### UPEC infection assay

Plate preparation: For the infection assays, OP50 and ECO85 were streaked directly from the frozen stock onto sheep-blood agar (Oxoid / Thermo Fisher, Waltham, MA). This passage over blood agar was critical for maximum pathogenicity in *C. elegans.* Overnight cultures were then grown in standard LB liquid at 37 °C for no more than 18 h. Next, 0.480 mL aliquots of this culture were spread on 6 cm petri plates containing 10 mL of high peptone (HP) NGM, i.e. 3.5 g/L tryptone/peptone from casein (Roth, Karsruhe, Germany) (compared to 2.5 g/L for standard NGM). Higher pathogenicity was obtained using tryptone/peptone from casein compared to Bacto-Peptone (BD Biosciences, Franklin Lakes, NJ). The plates were incubated overnight at 37 °C for 18-24h. OP50 and ECO85 were treated identically to eliminate strain-dependent uncontrolled effects due to growth on richer medium. Worm preparation: Embryos were extracted from gravid young adult using sodium hypochlorite treatment and hatched overnight in M9 buffer with agitation. The newly hatched larvae were transferred to standard NGM plates seeded with OP50 within 18 h of bleaching. These worms were then incubated at 20 °C until they reached the L4 larval stage (48 h). The L4 worms were then transferred to the ECO85 infection plates or OP50 mock-infection plates and incubated at 20 °C. Worms were scored for survival at 24 h intervals and death was defined as failure to respond to touch. Due to rapid declines in cuticle integrity, worms were not manipulated, i.e. transferred or moved, after the third day of infection tissue damage and premature death could result.

### Health parameters

Pharyngeal pumping was performed as described in (Bansal et al., 2015). The sensitivity of this assay was remarkable as temperature differences of only 2 to 3 °C could obscure the results; thus, the experiment was performed at precisely 20°C. Pharyngeal pumps were counted for 30 s for each worm. Because of the smaller changes in the pumping rate after injection of pure DNA, worms were monitored individually over three days. The pharyngeal pumping rates were measured for 30 sec intervals in triplicate for each worm with each measurement separated by at least 30 min to minimize behavioural effects or influences from other unrecognized variables. For crawling speed, movies of worms were produced using a Zeiss Axio Zoom V16 microscope. The crawling speed was then calculated using the WormLab software package (mbf Bioscience, Williston, VT). For fecundity: starting from day 1 of adulthood, worms were transferred in groups of three to fresh plates daily for 5 days. The total number of eggs produced was determined by counting both the number of eggs remaining on the plate, as well as the newly hatched progeny from eggs laid earlier. The surface area of the pharyngeal bulbs was performed by in Fiji (Schindelin et al., 2012) by drawing a circle around the pharyngeal metacorpus and posterior bulb to establish to regions of interest (ROIs) and then using the “Measure” function to determine their surface areas. The surface areas of the two ROIs in each worm were added, and the statistical analysis was performed on the combined values determined for each worm. The differential interference contrast (DIC) images of the pharynxes were collected using an AxioImager.M2 (Zeiss) and Zen Pro 2012 software (Zeiss). See also Supplemental Figure 1 for further description.

### In vivo staining

Nile Red powder (Sigma) was dissolved in acetone at a concentration 500 μg/ml. For treatment, this stock solution was diluted in 1X phosphate buffered saline (PBS) and pipetted onto NGM plates that were plates previously seeded with either OP50 or ECO85 (prepared as described in the UPEC infection experiment) to a final concentration of 0.05 μg/ml. Worms were transferred to these plates as L4 larvae (as in the UPEC infection experiment). Nile Red staining was monitored was monitored by fluorescence microscopy using AxioImager.M2 (Zeiss) and Zen Pro 2012 software (Zeiss). Acridine orange (AO) solution (Sigma) was diluted to a final concentration of 50 μg/mL in M9 buffer. Worms were picked from plates into M9 buffer and rinsed three times and then transferred into the AO solution. Following a 2 h incubation at 20 °C, the worms were transferred to NGM plates seeded with OP50 for 3 h for “destaining.” The AO staining was then monitored using AxioImager.M2 (Zeiss) and Zen Pro 2012 software (Zeiss).

### Immunofluorescence and confocal imaging

After 72 h of feeding on ECO85, N2 and *nuc-1(e1392)* worms were picked into 1 mL of M9 buffer and rinsed to remove excess bacteria. The intestines were then isolated vie decapitation in egg salts buffer with 0.2% Tween 20 (Roth). Tissue was fixed for 5 minutes in 3.7% formaldehyde (Roth) before being freeze cracked (Strome and Wood, 1982). The fixed tissues were treated with ice cold 100% methanol for 10 min. Following rinsing in PBS + 1% Triton X-100 (Roth) (PBSTX) and PBS + 0.1% Tween 20 (PBST) the samples were blocked for 1 h with normal donkey serum (NDS) (Sigma) diluted 1:10 in PBST before staining with antibody raised in mouse against α-tubulin (Sigma T6199). Secondary antibody was donkey Alexa Fluor 488-conjugated anti-mouse (Life Technologies / Thermo Fisher, Waltham, MA). Both were diluted 1:1000 in NDS. Samples were mounted in DAPI Fluoromount-G (SouthernBiotech, Birmingham, AL). Confocal imaging was performed using a Zeiss Meta 510 laser scanning confocal system and Zen 2009 software. Intracellular DAPI-stained inclusions were scored by centering 50 μm^2^ windows on nuclei of intestinal cells. The number of DAPI^+^ inclusions in the cytosol of the intestinal cells (i.e. in proximity to the α-tubulin signal) was counted over the entire thickness of the intestinal cells.

### Tunicamycin treatment

Tunicamycin (Sigma) dissolved in DMSO was diluted to a final concentration of 0.5 μg/mL. For the treatment, freshly hatched L1 larvae were grown on normal NGM plates seeded with OP50 (no tunicamycin) until the L4 stage. Infections were carried out as described above, except that the HP plates contained 0.5 μg/mL tunicamycin or an equal concentration of DMSO. For injection experiments, worms were grown on OP50-seeded NGM plates until they were injected on the first day of adulthood (as described above). After the injections, the worms were transferred to new OP50-seeded NGM plates containing the same concentration of tunicamycin. Imaging of live worms was performed using a Zeiss Axio ZoomV.16 microscope.

### RT-qPCR

Primers for assessment of UPR^ER^ activation were published earlier (Richardson et al., 2010). Other primers used:

*lys-7* GTCTCCAGAGCCAGACAATCC; CCAGTGACTCCACCGCTGTA

C14C6.5 CGGATACTGTGGCTCC; TATTTGCAATAACCGCATGT

*dct-17* GATGACTTCACATGTCCT; AGACATCGTACACTGCT

*T19D12.3* CTCACACGGATTATTACGG; AATAAGACCATCGGAAACTG

Following total RNA extraction using the RNeasy mini-kit (Qiagen), complementary DNA was production using the Superscript II protocol (Invitrogen). Quantitative real-time (RT-qPCR) was carried out using Biorad CFX96 real-time PCR machines with SYBR Green I for amplification quantification (Sigma) and Platinum *Taq* polymerase (Invitrogen/ Thermo Fisher, Waltham, MA). That specific products were generated was confirmed via melting curve analysis. For data analysis, the comparative C_T_(2^−ΔΔc^_T_) method (Livak and Schmittgen, 2001) was used, where 2^−ΔΔc^_T_ = [(C_T_gene of interest – C_T_internal control) sample A) – (C_T_gene of interest – C_T_internal control sample B)]. Expression levels were normalized using the actin (*act-1*) and tubulin (*tbg-1*) genes as internal controls and standard deviations were calculated as in (Schmittgen and Livak, 2008).

### N-acetylglucosamine treatment

*N*-acetylglucosamine (GlcNAc) (Sigma) dissolved in water at a stock concentration of 500 mM. Worms were grown on OP50-seeded NGM plates until they were injected on the first day of adulthood as described above. After the injections, the worms were transferred to new OP50-seeded NGM plates containing a final concentration of 10 mM GlcNAc. Pumping was scored daily as described above.

### Transmission electron microscopy

Sample preparation was performed as in (Hall et al., 2012). Briefly, worms were removed from food and fixed in 0.8% glutaraldehyde in 0.1 M sodium cacodylate buffer, pH 7.3. Worms were then treated with 0.8% OsO_4_ in the same buffer. Worms were washed to remove glutaraldehyde and incubated in 2% OsO_4_ in 0.1 M sodium cacodylate buffer for 24 h at 4 °C. Worms were then embedded in 2% agarose and dehydrated via treatment with increasing concentrations of ethanol (50% - 100%) followed by embedding in Epon before preparation of thin slices for imaging. Microscopy was performed using a Zeiss EM109 electron microscope (Zeiss, Oberkochen, Germany).

